# Flexibility and mobility of SARS-CoV-2-related protein structures

**DOI:** 10.1101/2020.07.12.199364

**Authors:** Rudolf A. Römer, Navodya S. Römer, A. Katrine Wallis

**Affiliations:** CY Advanced Studies and LPTM (UMR8089 of CNRS), CY Cergy-Paris Université, F-95302 Cergy-Pontoise, France; Department of Physics, University of Warwick, Coventry, CV4 7AL, United Kingdom; School of Life Sciences, University of Lincoln, Brayford Pool Campus, Lincoln, LN6 7TS, United Kingdom; School of Life Sciences, University of Warwick, Coventry, CV4 7AL, United Kingdom

## Abstract

The worldwide CoVid-19 pandemic has led to an unprecedented push across the whole of the scientific community to develop a potent antiviral drug and vaccine as soon as possible. Existing academic, governmental and industrial institutions and companies have engaged in large-scale screening of existing drugs, in vitro, in vivo and in silico. Here, we are using in silico modelling of SARS-CoV-2 drug targets, i.e. SARS-CoV-2 protein structures as deposited on the Protein Databank (PDB). We study their flexibility, rigidity and mobility, an important first step in trying to ascertain their dynamics for further drug-related docking studies. We are using a recent protein flexibility modelling approach, combining protein structural rigidity with possible motion consistent with chemical bonds and sterics. For example, for the SARS-CoV-2 spike protein in the open configuration, our method identifies a possible further opening and closing of the S1 subunit through movement of S^B^ domain. With full structural information of this process available, docking studies with possible drug structures are then possible in silico. In our study, we present full results for the more than 200 thus far published SARS-CoV-2-related protein structures in the PDB.

## Introduction

At the end of 2019 a cluster of pneumonia cases was discovered in Wuhan city in China, which turned out to be caused by a novel coronavirus, SARS-CoV-2.^1^ Since then the virus has spread around the world and currently has caused over 10 million infections with more than 500,000 deaths worldwide (July 1st).^2^ SARS-CoV-2 is the seventh coronavirus identified that causes human disease. Four of these viruses cause infections similar to the common cold and three, SARS-CoV, MERS-CoV and SARS-CoV-2 cause infections with high mortality rates.^3^ As well as affecting humans, coronaviruses also cause a number of infections in farm animals and pets,^4^ and the risk of future cross-species transmission, especially from bats, has the potential to cause future pandemics.^5^ Thus, there is an urgent need to develop drugs to treat infections and a vaccine to prevent this disease. Some success has already been achieved for SARS-CoV-2 with dexamethasone reducing mortality in hospitalised patients^6^ and a number of vaccine trials are currently ongoing.

The viral spike protein is of particular interest from a drug- and vaccine-development perspective due to its involvement in recognition and fusion of the virus with its host cell. The spike protein is a heavily glycosylated homotrimer, anchored in the viral membrane. It projects from the membrane giving the virus its characteristic crown-like shape.^3^ The ectodomain of each monomer consists of an N-terminal subunit, S1, comprising two domains, S^A^ and S^B^, followed by an S2 subunit forming a stalk-like structure. Each monomer has a single membrane-spanning segment and a short C-terminal cytoplasmic tail.^7^ The S1 is involved in recognition of the human receptor, ACE2. This subunit has a closed or down configuration where all the domains pack together with their partners from adjacent polypeptides.^8, 9^ However, in order for recognition and binding to ACE2 to take place, one of the three S^B^ domains dissociates from its partners and tilts upwards into the open or up configuration.^8, 9^ Binding of ACE2 to the open conformation leads to proteolytic cleavage of the spike polypeptide between S1 and S2.^7^ S2 then promotes fusion with the host cell membrane leading to viral entry.^10^ Drugs that target the spike protein thus have the potential to prevent infection of host cells.

The main protease of the virus (M^pro^) is another important drug target. M^pro^ is responsible for much of the proteolytic cleavage required to obtain the functional proteins needed for viral replication and transcription. These proteins are synthesised in the form of polyproteins which are cleaved to obtain the mature proteins.^11^ M^pro^ is active in its dimeric form but the SARS-CoV M^pro^ is found as a mixture of monomer and dimers in solution.^12^ SARS-CoV M^pro^ has been crystallised in different, pH-dependent conformations suggesting flexibility, particularly around the active site. MD simulations support this flexibility.^13^ The protease is highly conserved in all coronaviruses so targeting either dimerization or enzymatic activity may give rise to drugs that can target multiple coronaviruses, known and yet unknown.^14, 15^

Since the discovery of SARS-CoV-2, a plethora of structures have been determined including M^pro16^ and the ectodomain of the spike protein^8, 9^ as well as other potential drug and vaccine targets. These structures provide the opportunity for rational drug design using computational biology to identify candidates and optimise lead compounds. However, crystal structures only provide a static picture of proteins, whereas proteins are dynamic and this property is often important in drug development. For example, agonists and antagonists often bind different conformations of G coupled-protein receptors.^17^. Flexibility also affects thermodynamic properties of drug binding,^18^, yet the ability to assess flexibility is often hampered by the long computational times needed for MD simulations.

We use a recent protein flexibility modeling approach,^19^ combining methods for deconstructing a protein structure into a network of rigid and flexible units (First)^20^ with a method that explores the elastic modes of motion of this network,^21–25^ and a geometric modeling of flexible motion (Froda).^26, 27^ The usefulness of this approach has recently been shown in a variety of systems.^28–32^ Methods similar in spirit, exploring flexible motion using geometric simulations biased along “easy” directions, have also been implemented using Frodan^33^ and NMSim.^34^ We have performed our analysis through multiple conformational steps starting from the crystal structures of SARS-CoV-2-related proteins as currently deposited in the PDB. This results in a comprehensive overview of rigidity and the possible flexible motion trajectories of each protein. We emphasize that these trajectories do not represent actual stochastic motion in a thermal bath as a function of time, but rather the possibility of motion along a set of most relevant low-frequency elastic modes. Each trajectory leads to a gradual shift of the protein from the starting structure and this shift may reach an asymptote, where no further motion is possible along the initial vector, as a result of steric constraints.^31^ Energies associated with such a trajectory for bonds, angles, electrostatics, and so forth, have be estimated in previous studies of other proteins and shown to be consistent and physically plausible. Computing times for the method vary with the size of the proteins, but range from minutes to a few hours and are certainly much faster than thermodynamically equilibrated MD simulations. The approach hence offers to possibility of large-scale screening of protein mobilities.

## Results

### Protein selection

We have downloaded protein structure files as deposited on the Protein Data Bank,^35^ including all PDB codes that came up when using “SARS-CoV-2” and “Covid-19” as search terms, as well as minor variations in spelling. In our first search of April 18th, 2020, this gave 133 protein structures. A second and third search on May 20th and 29th, 2020, respectively, resulted in a total of 219 structures. In addition we included a few protein structures from links provided in selected publications to give a grand total of 223 protein structure included in this study. Many of the structures found, as outlined above, have been deposited on the PDB in dimer or trimer forms. Hence one has the choice to study the rigidity and motion of the monomer or the dimer/trimer. Clearly, the computational effort for a dimer/trimer is much larger than for a monomer since in addition to the intra-monomer bond network, also inter-monomer bond need to be taken into account. Furthermore, it is not necessarily clear whether the possible motion of a monomer or dimer should be computed to have the most biological relevance. We have computed the motion of full dimer and trimer structures only for a few selected and biologically most relevant such structures while the default results concentrated on the monomers. Nevertheless, we wish to emphasize that when results for a certain monomer exist, it should nearly always be possible to also obtain their motion in dimer/trimer configuration. For some protein structures, we have found that steric clashes were present in the PDB structures that made a flexibility and sometimes even just the rigidity analysis impossible. Usually, this is due to a low crystal resolution. A list of all current protein PDB IDs included in this work is given in Table S1.^1^

### Rigidity and Flexibility

In Figs. 1 and 2 we show examples of different rigidity patterns that emerge from the First analysis. In line with previous studies comparing these rigidity patterns across various protein families,^19^ we find that they can be classified into about four types. In Fig. 1(a) we see that for the crystal structure of SARS-CoV-2 nucleocapsid protein N-terminal RNA binding domain (PDB:6m3m), the largest rigid cluster in the pristine structure, i.e. at *E*_cut_ = 0, largely remains rigid through the dilution process of consecutively lowering *E*_cut_ values. When bonds are opened the rigid cluster shrinks but the newly independent parts are flexible and not part of any new large rigid cluster themselves. Only very small parts of the protein chain break to form their own independent rigid structures before at a certain *E*_cut_ the whole protein is essentially flexible. We shall denote such a behaviour as *brick*-like. In contrast, in Fig. 1(b) we observe that for chain A of the co-factor complex of NSP7 and the C-terminal domain of NSP8, which forms part of the RNA synthesis machinery from SARS CoV-2 (PDB:6wiq), already the crystal structure has fallen into three independent rigid structures. Opening bonds, we find that the largest rigid cluster breaks into 2 rigid structures.

**Figure 1.**
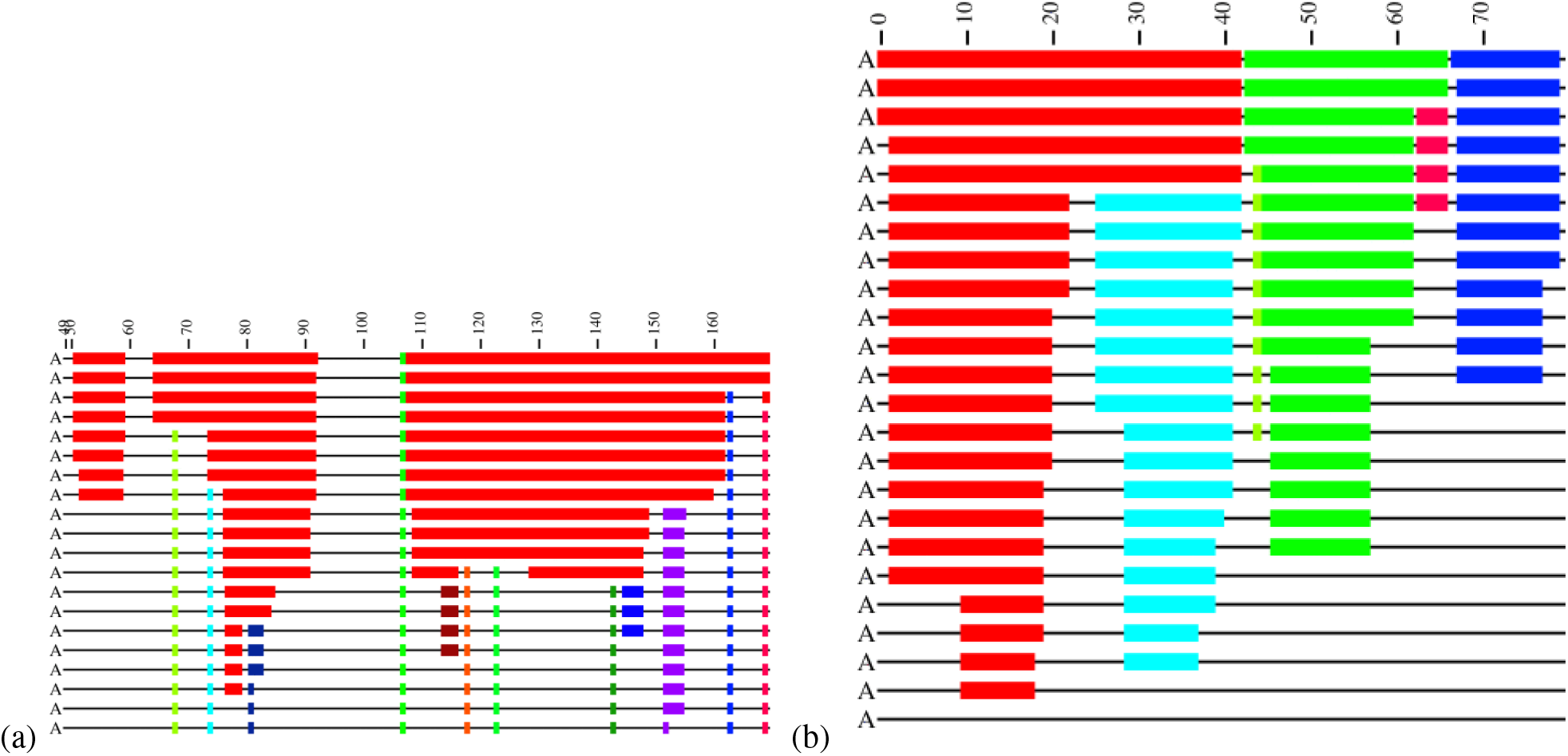
Examples of rigid cluster decompositions for different SARS-CoV-2 protein structures with (a) showing the crystal structure of SARS-CoV-2 nucleocapsid protein N-terminal RNA binding domain (PDB:6m3m) while (b) gives chain A of the co-factor complex of NSP7 and the C-terminal domain of NSP8 from SARS CoV-2 (PDB:6wiq). Different rigid clusters of the polypeptide chain appear as identically coloured blocks along the protein chain with each *C*_*α*_ labelled from its N-terminal at 1 to its C-terminal. When the energy cutoff *E*_cut_ decreases (left-most column, downward direction towards larger, negative *E*_cut_ magnitudes), rigid clusters break up and more of the chain becomes flexible. The colour coding shows which atoms belong to which rigid cluster. Flexible regions appear as black thin lines. The second column on the left indicates the mean number ⟨*r*⟩ of bonded neighbours per atom as *E*_cut_ changes. We note that the *E*_cut_ scale is not linear since new rows are only added when the rigidity of a structure has changed.

**Figure 2.**
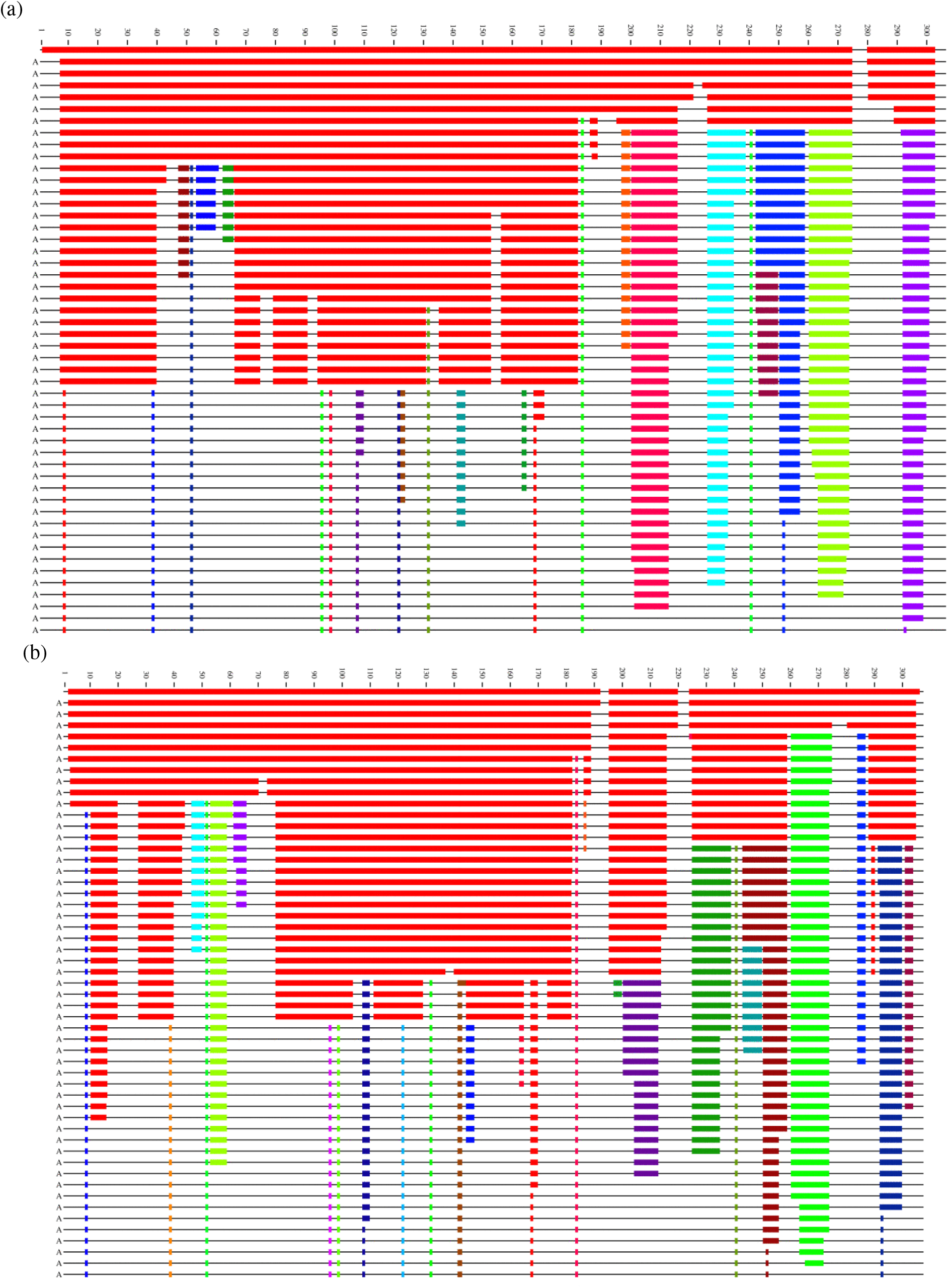
Examples of rigid cluster decompositions for different SARS-CoV-2 protein structures. (a) show the monomer of crystal structure of COVID-19 main protease in apo form (PDB:6m03) and (b) is chain A of complex resulting from the reaction between the SARS-CoV main protease and a carbamate (PDB:6y7m). Different rigid clusters of the polypeptide chain appear as identically coloured blocks along the protein chain with each *C*_*α*_ labelled from its N-terminal at 1 to its C-terminal. When the energy cutoff *E*_cut_ decreases (left-most column, downward direction towards larger, more negative *E*_cut_ magnitudes), rigid clusters break up and more of the chain becomes flexible. The colour coding shows which atoms belong to which rigid cluster. Flexible regions appear as black thin lines. The second column on the left indicates the mean number ⟨*r*⟩ of bonded neighbours per atom as *E*_cut_ changes.

Then these now four *domains*^2^ retain their character upon opening more and more bonds until they simply dissolve into full flexibility.

While brick- and domain-like behaviour is found only occasionally, more prevalent are two further types of rigidity dissolution. In Fig. 2(a) we see that for, e.g. the monomer of crystal structure of COVID-19 main protease (M^pro^) in apo form (PDB:6m03), the rigid cluster dominating the crystal structure quickly falls apart upon change in *E*_cut_ with five newly formed independent rigid clusters emerging towards to N-terminal of the protein chain. These new clusters remain stable to the opening of further bonds, even when the remnants of the original rigid cluster has become fully flexible. Such a behaviour relies on a certain critical *E*_cut_ value to dominate the rigidity dissolution and is reminiscent of so-called first-order phase transitions in statistical physics. Hence this rigidity-type is usually denoted as *1st order*.^19^ The behaviour seen in 2(b) for the crystal structure of the complex resulting from the reaction between the SARS-CoV main protease and tert-butyl (1-((S)-3-cyclohexyl-1-(((S)-4-(cyclopropylamino)-3,4-dioxo-1-((S)-2-oxopyrrolidin-3-yl)butan-2-yl)amino)-1-oxopropan-2-yl)-2-oxo-1,2-dihydropyridin-3-yl)carbamate (PDB:6y7m) is markedly different. Here, there are many values of *E*_cut_ where large parts of the original rigid structure break off one after another so that towards the end of the bond-opening process, the original cluster is still present, but no longer dominates the rigidity pattern. This behaviour is called *2nd order*.

In Table S1, we have indicated the classification for each protein structure into the four classes. Obviously, this classification is not perfect and there are also sometimes intermediate rigidity patterns. Nevertheless, this rigidity classification already provides a first insight into the possible flexibility and range of motion for each structure. It should be clear, that a brick-like rigidity should show the least flexibility until it dissolves completely. On the other hand, for domain-like structures, one can expect possible intra-domain motion while inter-domain motion might be harder to spot. Similarly, for a 1st-order rigidity, one would expect little dynamic mobility until the “transition” value for *E*_cut_ has been reached, although high levels of flexibility should be possible afterwards. Last, a protein with 2nd-order rigidity should have the most complex behaviour in terms of flexibility since new possible mobility can be expected throughout the range of *E*_cut_ values.

### Protein Mobility

For each value of *E*_cut_, the First analysis has provided us with a map of rigid and flexible regions in a given crystal structure. We can now translate this into propensity of motion by allowing the flexible parts to move perturbatively, subject to full bonding and steric constraints (see Methods section). Moving along directions proposed by an elastic normal model analysis of the crystal structure, we can therefore construct possible motion trajectories that are fully consistent with the bond network and steric constraints. Each trajectory corresponds to one such normal mode, denoted *m*_7_ up to *m*_12_ for the first low-frequency non-trivial such modes, as well as the chosen *E*_cut_ value. Generally, a larger value of |*E*_cut_| implies less rigidity and results in larger scale flexible motion. In Fig. 3 we give examples of such motion trajectories. Fig. 3(a) shows a monomer of the SARS-CoV-2 spike glycoprotein (closed state) (PDB code: 6vxx^9^). We can see that there is a good range of motion from the crystal structure when following the normal mode modes into either positive or negative changes along the mode vector. Fig. 3(b) shows the motion for the dimer structure of the SARS-CoV-2 main protease (PDB code: 6lu7, in complex with inhibitor N3)^16^. Again, one can see considerable motion, although due to the complexity of the structure, it is difficult to distinguish individual movement patterns from such a frozen image. Much better insight can be gained when watching for full range of motion as a movie. We have therefore included movies for all figures as supplemental materials. We also have made a dedicated web download page^36^ where movies for the complete set of PDB codes as given in Table S1 are publicly available. The movies are being offered for various modes *m*_7_ to *m*_12_ as well as *E*_cut_ values, at least containing results for *E*_cut_ = 1 kcal/mol, 2 kcal/mol and 3 kcal/mol, respectively. In addition, the site includes the rigidity resolutions discussed above and all intermediate structures needed to make the movies. This allows for the calculation of relative distances and other such structural measures along each motion trajectory as desired. For a detailed analysis of the SARS-CoV-2 main protease with similar methods, we refer the reader to Ref. 26.

**Figure 3.**
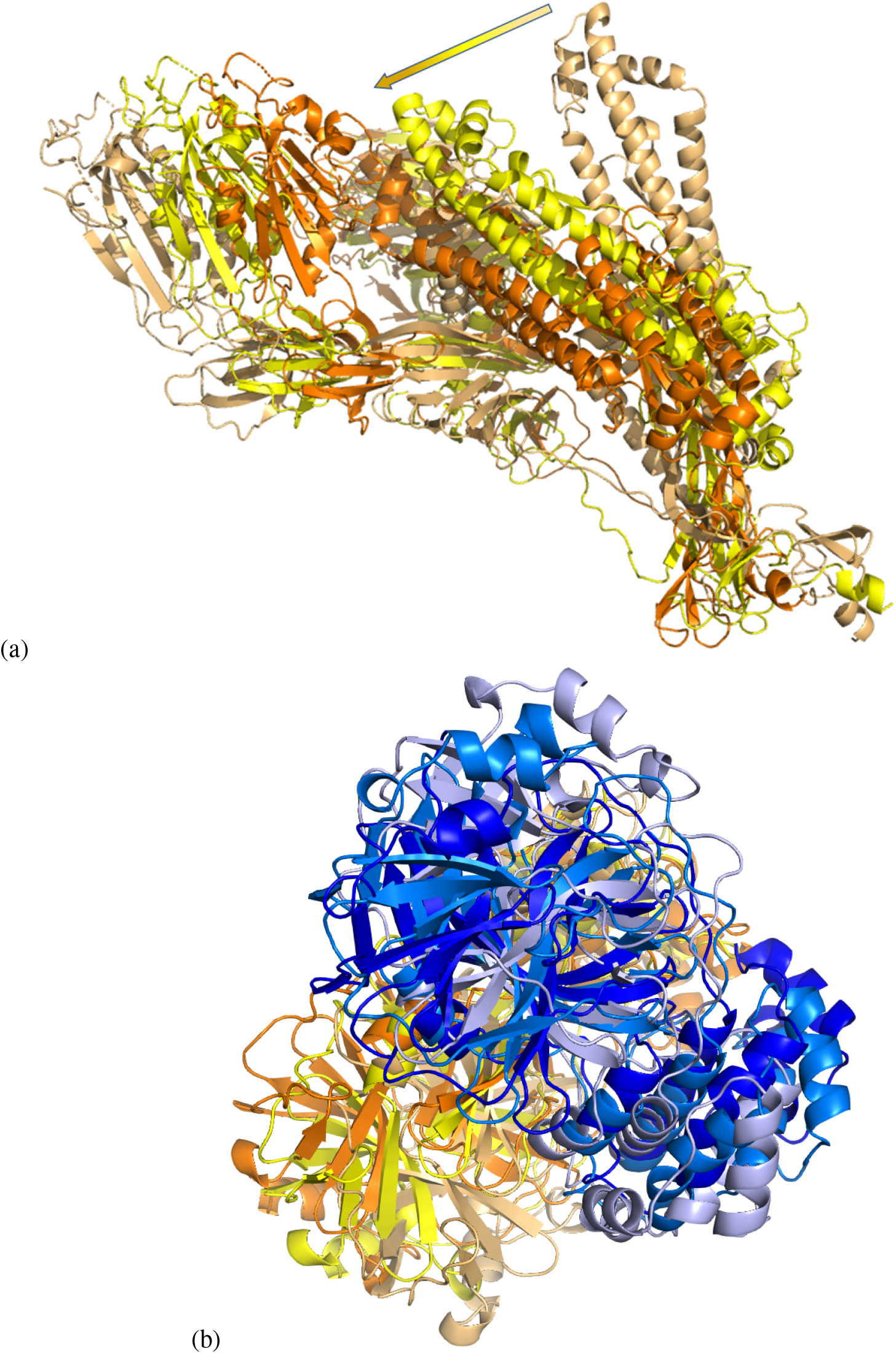
Examples of flexible motion for (a) mode *m*_7_ of the SARS-CoV-2 spike ecto domain (6vxx) monomer at *E*_cut_ = 0.5 kcal/mol with chain A denoted by yellow and its extreme structural positions given as light orange/orange. In panel (b), we show *m*_7_ for the dimer of main protease (PDB: 6lu7) at *E*_cut_ = 2 kcal/mol with chain A denoted again by yellow and extremal structures colored as in (a); chain B is indicated in blue with extremal positions in light and dark blue. The arrow in (a) indicates the range of movement for the dominant *α*-helix structure in the spike ecto domain, from negative to positive movement.

### SARS-CoV-2 spike ecto domain structures

As discussed above, the trans-membrane spike glycoprotein mediates entry into host cells.^8, 9^ As such, it is a “main target for neutralizing antibodies upon infection and the focus of therapeutic and vaccine design”.^9^ It forms homotrimers protruding from the viral surface.^3^ Up to the end of May 2020, 3 structures of the trans-membrane spike glycoprotein have been deposited in the PDB. These transmission electron cryomicroscopy (cryo-EM) studies have led to structures with PDB codes 6vsb^8^, 6vxx^9^ and 6vyb^9^ now being available. With RMS resolution of 3.5 Å, 6vsb has a slightly lower resolution than 6vxx at 2.8 Å and 6byv at 3.2 Å. In the following, we shall discuss the resulting rigidity and flexibility properties of these three structures in their full trimer form. Results for individual monomer rigidities, as given in Fig. 3(a), are also available at Ref. 36.

#### Prefusion 2019-nCoV spike glycoprotein with a single receptor-binding domain open

The rigidity pattern of the homotrimer for this structure (PDB:6vsb, 440.69 kDa, 22854 atoms, 2905 residues) is dominated by a large rigid cluster encompassing most of the trimer structure except for a region from roughly residue 330 to residue 525.^3^ This indeed corresponds to the S^B^ domain of S1 in each monomer. In the trimer configuration, it is known that one of the S^B^ domains can change from the closed to an open configuration^8^ At |*E*_cut_| = 0.016 kcal/mol, the 6vsb structure of the trimer breaks into many different rigid parts, but the original large cluster remains a dominating feature across the rigidity dilution plot. A motion analysis analysis does not compute but breaks with bad sterics, apparently due to the comparably low resolution of the crystal structure.

#### Mobility of the SARS-CoV-2 spike glycoprotein (closed state)

For the closed state structure (6vxx, 438.53 kDa, Atom Count: 23694 atoms, 2916 residues) of the SARS-CoV-2 spike glycoprotein, the rigidity pattern is again “brick”-like and the whole of the crystal structure is part of a single cluster. At |*E*_cut_| = 0.022 kcal/mol, there is a first order break of the large rigid cluster into dozens of smaller rigid units. Nevertheless, the original cluster retains a good presence in each chain of the trimer. In terms of motion, we are now able to produce motion studies for the full set of modes *m*_7_ to *m*_12_ and various *E*_cut_’s. In Fig. 4 we show motion results, using again the structures for the extreme ranges of the motion as in Fig. 3. In addition, we are showing side, c.p. Fig. 4(a) and top, c.p. Fig. 4(b), views similar to Ref. 9. Looking at the whole range of modes and *E*_cut_’s computed, we find that motion is very reminiscent of the vibrational excitations of a rigid cone or cylinder. There is twist motion around the central axis, bending of the trimer along the central axis, relative twist motion of 2 chains relative to the remaining chain (c.p. Fig. 4(b)), etc. Already at *m*_12_ (with |*E*_cut_| = 2 kcal/mol), large scale motion has stopped and one only observes smaller scale motion is flexible parts of the trimer chains. Overall, this behaviour is very consistent with the elastic behaviour of a “closed” structure^9^ similar to a cone or cylinder.

**Figure 4.**
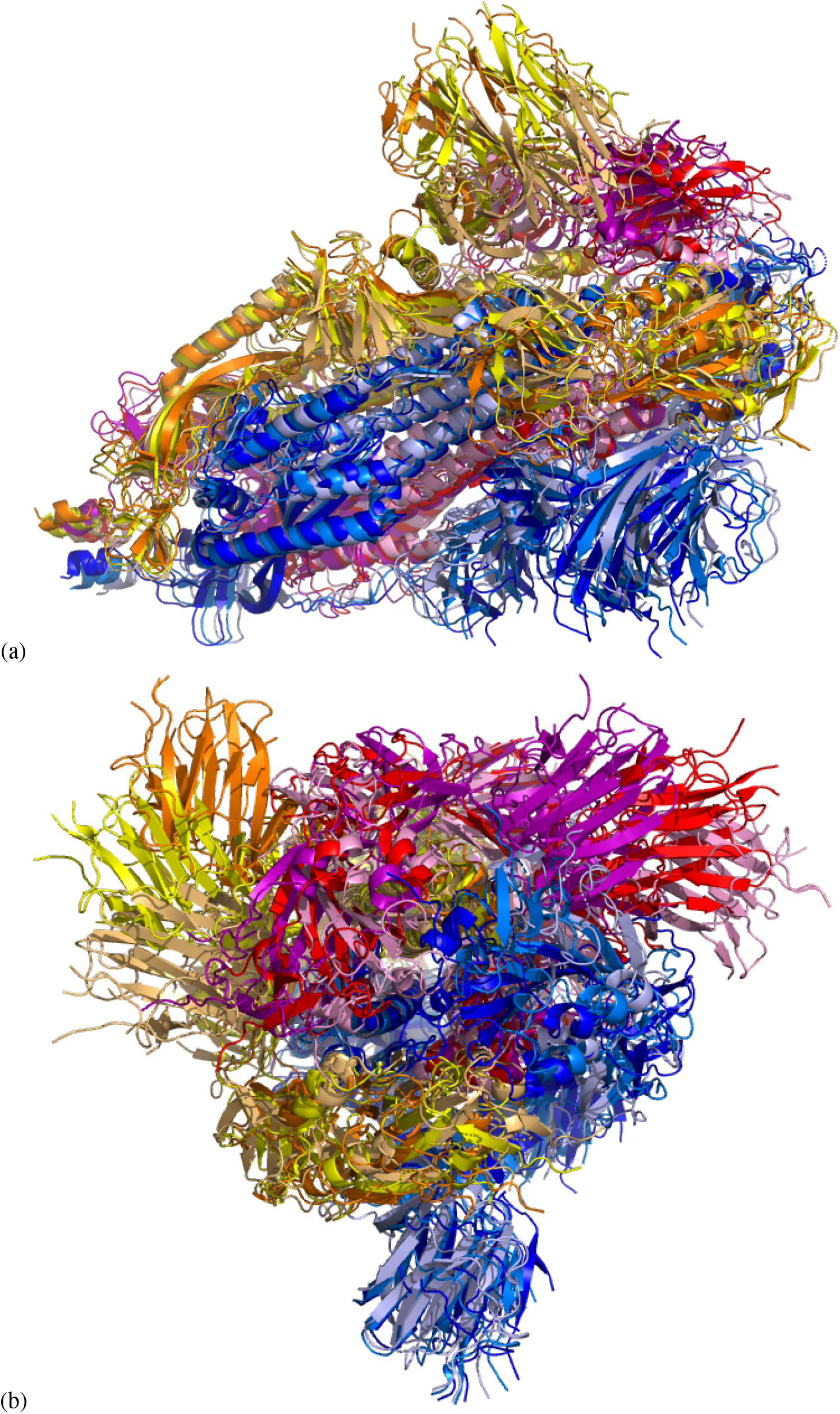
Two views of possible motion in the spike ecto domain in closed configuration (6vxx)^9^ with a view of *m*_7_ motion at *E*_cut_ = 2 kcal/mol (a) from the side and (b) from the top for the whole trimer. The secondary protein structure is highlighted by the chosen “cartoon” representation. Colors yellow, blue and red denote chains A, B, and C while color combinations light orange/orange, pink/purple and light blue/dark blue show the extreme structural positions for the movements along the normal mode *m*_7_ at *E*_cut_ = 2 kcal/mol.

#### Mobility of the SARS-CoV-2 spike glycoprotein (open state)

We now turn to the last SARS-CoV-2 spike ecto domain structure (6vyb, 437.65 kDa, 22365 atoms, 2875 residues).^9^ This structure is in the open configuration similar to the conformation seen in 6vsb.^8^ The structure is shown in Fig. 5, again in side and top view. From the rigidity plots we find, in addition to the largest rigid cluster, already for the crystal structure at |*E*_cut_| = 0 kcal/mol a second rigid cluster, also roughly spanning from residue 330 to 525. Again, this region identifies the S^B^ domain as in the 6vsb structure. Compared to the closed state (6vxx), we see that this cluster has more internal structure, i.e. consists of more flexible parts in the S^B^ region from 330 to 525 and also seems to fall apart upon further changing *E*_cut_. This suggests that it is more flexible as a whole. Upon motion simulation, we observe a very high mobility in that prominent S^B^ subdomain. Starting from the crystal structure, we find for |*E*_cut_| = 1 kcal/mol a clear further opening towards a negative FRODA mode, *m*_7_, while in the other, positive direction of *m*_7_, the structure can again close the trimer. The distance range of the motion can be expressed as follow: the distance from residue 501 in the middle of the central *β* -sheet of the S^B^ to the most opened conformation is 57 Å while distance to the same residue in the most closed conformation is 28 Å. The distance from open to closed is 69 Å. Hence the motion simulation adds additional insight into the distinction between the open (6vyb) and the closed (6vxx) structures while also showing that a transition from open to closed is indeed possible. As stated in Ref. 9, this interplay of closing and opening is expected to be central to the viral entry into the human cell.

**Figure 5.**
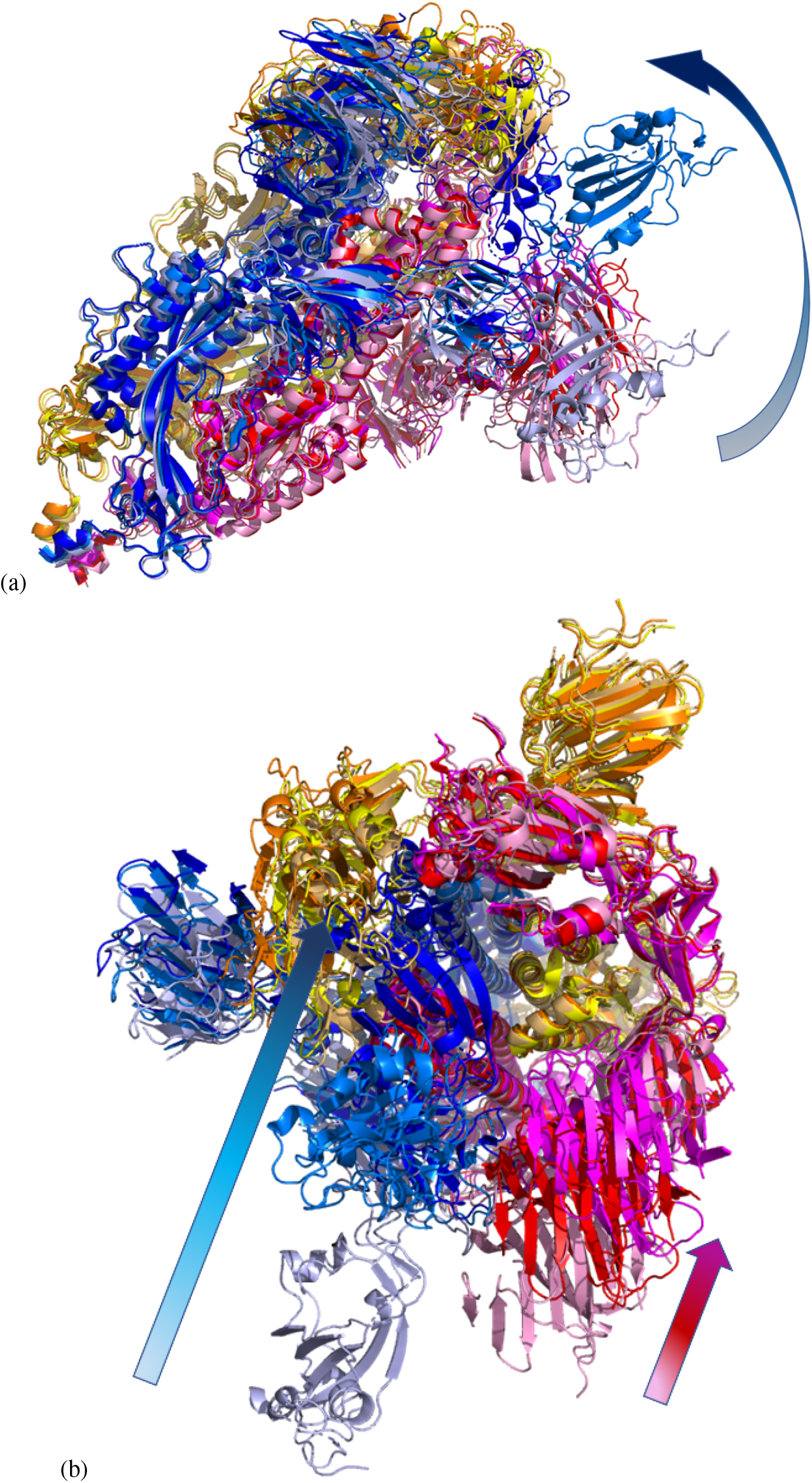
Possible motion along mode *m*_7_ at *E*_cut_ = 1 kcal/mol with (a) side and (b) top view of the open spike ecto domain (6vyb)^9^. Colors are chosen identical to Fig. 4. As in Fig. 3, the arrows in each panel show the range of motion for chain B (blue shades) in (a+b) and also for chain C (reds) in (b).

## Discussion

SARS-CoV-2 infectivity is dependent on binding of the spike protein to ACE2. This binding is only possible when the spike protein is in the open conformation. Structures of both the open and closed conformation have already been determined and the flexibility of the S^B^ domain inferred from these static structures^8, 9^ . The spike trimer consists of almost 3000 amino acids and hence is not an easy target for dynamics simulations due to its size. In the open structures (6vsb and 6vyb) the S^B^ domain is clearly identifiable in the rigidity analysis as a separate cluster to the rest of the trimer. This shows that this domain has increased flexibility. We can clearly observe the hinge movement of the S^B^ domain in the open configuration (6vyb) with the S^B^ domain moving back into the closed configuration. The range of movement from the most opened to the most closed conformation can be measured to be quite large and the flexibility within the S^B^ domain itself during the hinge movement is also seen to be considerable. All these findings suggest that the S^B^ domain of the spike protein has the necessary flexibility to attach itself readily to ACE2. However, when starting from the closed structure (6vxx), we do not see an opening. This suggests possibly stronger bonds and steric constraints which need to be overcome before the structure is able to open up. Nevertheless, to our knowledge this is the first time the hinge motion of the S^B^ domain has been predicted solely based on the dynamics of a structure. In principle, the full structural information provided in our download site^36^ can be used to perform possible docking studies with ligands of different sizes. These intermediate conformations can also be used as convenient starting points for further MD-based dynamics studies to evaluate thermodynamic properties of the structures. This shows the power of FIRST and FRODA to complement structure determinations and MD simulations to make valid predictions about the dynamics of proteins. Similarly detailed structural analyses and further MD work is also possible for the other more than 200 structures included in this work. The classification of the protein structures into four types of dynamics provides information about the level of flexibility of a protein which is relevant to drug discovery. For more detailed information rigidity dilution plots and motion movies are available for download at Ref. 36. In addition, the code underlying our study is accessible at Ref. 37 and implemented in a free web server at Ref. 38.

## Methods

### Rigidity analysis

We start the rigidity, flexibility and mobility modelling in each case with a given protein crystal structure file in PDB *.pdb* format. Hydrogen atoms absent from the PDB X-ray crystal structures are added using the software Reduce^39^. Alternate conformations that might be present in the protein structure file are removed if needed and the hydrogen atoms renumbered in PyMol.^40^ We find that for some protein structures the addition of hydrogen atoms is not possible without steric clashes. Consequently, identification of a viable structure and its continued analysis is not possible. These proteins are labelled in Table S1.

For the remaining proteins we produce the ‘rigidity dilution’ or rigid cluster decomposition (RCD) plot using First.^20^ The plots show the dependence of the protein rigidity on an energy cutoff parameter, *E*_cut_ < 0. It parametrizes a bonding range cutoff based on a Mayo potential^41, 42^ such that larger (negative) values of *E*_cut_ correspond to more open bonds, i.e. a smaller set of hydrogen bonds to be included in the rigidity analysis.

### Elastic network modes

We obtain the normal modes of motion using elastic network modelling (ENM)^43^ implemented in the ElNemo software.^23, 24^ This generates a set of elastic eigenmodes and associated eigenfrequencies for each protein. The low-frequency modes are expected to have the largest motion amplitudes and thus be most significant for large conformational changes. The six lowest-frequency modes (modes 1–6) are just trivial combinations of rigid-body translations and rotations of the entire protein. Here we consider the six lowest-frequency non-trivial modes, that is, modes 7–12 for each protein. We will denote these modes as *m*_7_, *m*_8_, …, *m*_12_.

### Mobility simulations

The modes are next used as starting direction for a geometric simulation, implemented in the Froda module^26^ within First. This explores the flexible motion available to a protein within a given pattern of rigidity and flexibility. Froda then reapplies bonding and steric constraints to produce an acceptable new conformation. Since the displacement from one conformation to the next is typically small, we record only every 50th conformation. The computation continues for typically several thousand conformations. A mode run is considered complete when no further projection along the mode eigenvector is possible (due to steric clashes or bonding constraints). This manifests itself in slow generation of new conformations.

We have performed Froda mobility simulation for each protein at several selected values of *E*_cut_. This allows us to study each protein at different stages of its bond network, roughly corresponding to different environmental conditions, such as different temperatures as well as different solution environments. In a previous publication, we discussed the criteria for a robust selection of *E*_cut_.^42^ Ideally, for each protein structure, a bespoke set of *E*_cut_ values should be found, with the RCD plots providing good guidance on which *E*_cut_ values to select. Clearly, for a large-scale study as presented here, this is not readily possible due to time constraints. Instead, we have chosen *E*_cut_ = −1 kcal/mol, −2 kcal/mol and −3 kcal/mol for each protein. These values have been used before in a multi-domain protein with 22 kDalton and shown to reproduce well the behaviour of (i) a mostly rigid protein at *E*_cut_ = −1 kcal/mol, (ii) a protein with large flexible substructures/domains at *E*_cut_ = −2 kcal/mol and (iii) a protein with mostly flexible parts connecting smaller sized rigid subunits at *E*_cut_ = −3 kcal/mol.^31, 42^ In addition, we have also performed the analysis at other values of *E*_cut_ when upon inspection of the RCD plots it was seen that the standard values (i) – (iii) would not be sufficient. The exact values used are given in Table S1.

We emphasize that these trajectories do not represent actual stochastic motion in a thermal bath as a function of time, but rather the *possibility of motion* along the most relevant elastic modes. Each trajectory leads to a gradual shift of the protein from the starting structure. This shift eventually reaches an asymptote, where no further motion is possible along the initial vector, as a result of steric constraints. Energies associated with such a trajectory for bonds, angles, electrostatics, and so forth, can be estimated and shown to be consistent and physically plausible.^31^

## Acknowledgements

This work received funding by the CY Initiative of Excellence (grant “Investissements d’Avenir” ANR-16-IDEX-0008) and developed during R.A.R.’s stay at the CY Advanced Studies, whose support is gratefully acknowledged. We thank Warwick’s Scientific Computing Research Technology Platform for computing time and support. Special thanks to overleaf.com for free premium access during Covid-19 lockdown. UK research data statement: Data accompanying this publication are available for download.^36^

## Author contributions statement

RAR conceived the study, NSR and RAR assembled Table S1 and identified *E*_cut_ values, RAR performed the computations and curated the data, AKW identified biologically most relevant structures. All authors wrote and reviewed the manuscript.

## Additional information

### Competing interests

The authors declare no competing interests.

## Footnotes

1 We expect to continue to increase the number of SARS-CoV-2-related proteins to be included in our study as more structures become available.

2 We use the term *domain* here to denote large regions of a protein connected in the same rigidity cluster. Usually, domains in a biological sense are also detected as domains in a rigidity sense as used here.^32^ However, this relationship is not necessarily always the case.

3 With nearly 3000 residues in the structure of the trimer, is it impossible to capture the behaviour in a single figure that at the same time would allow the reader to see enough detail. Nevertheless, we include the figure among the supplementary materials so that both an overview can be achieved as well as enough detail studied by using standard scalable postscript document viewers.

## Notes

https://www.warwick.ac.uk/flex-covid19-data

### Competing Interest Statement

The authors have declared no competing interest.

## References

1. Chan, J. F. W. et al. A familial cluster of pneumonia associated with the 2019 novel coronavirus indicating person-to-person transmission: a study of a family cluster. The Lancet 395, 514–523, DOI: 10.1016/S0140-6736(20)30154-9 (2020).

2. WHO Coronavirus Disease (COVID-19) Dashboard | WHO Coronavirus Disease (COVID-19) Dashboard.

3. Monto, A. S., Cowling, B. J. & Peiris, J. S. Coronaviruses. In Viral Infections of Humans: Epidemiology and Control, 199–223, DOI: 10.1007/978-1-4899-7448-8{_}10 (Springer US, 2014).

4. Tizard, I. R. Vaccination Against Coronaviruses In Domestic Animals. Vaccine 38, 5123–5130, DOI: 10.1016/j.vaccine.2020.06.026 (2020).

5. Letko, M., Seifert, S. N., Olival, K. J., Plowright, R. K. & Munster, V. J. Bat-borne virus diversity, spillover and emergence, DOI: 10.1038/s41579-020-0394-z (2020).

6. Horby, P. et al. Dexamethasone for COVID-19-Preliminary Report Effect of Dexamethasone in Hospitalized Patients with COVID-19-Preliminary Report RECOVERY Collaborative Group*. medRxiv 2020.06.22.20137273, DOI: 10.1101/2020.06.22.20137273 (2020).

7. Li, F. Structure, Function, and Evolution of Coronavirus Spike Proteins. Annu. Rev. Virol. 3, 237–261, DOI: 10.1146/annurev-virology-110615-042301 (2016).

8. Wrapp, D. et al. Cryo-EM structure of the 2019-nCoV spike in the prefusion conformation. Science 367, 1260–1263, DOI: 10.1126/science.abb2507 (2020).

9. Walls, A. C. et al. Structure, Function, and Antigenicity of the SARS-CoV-2 Spike Glycoprotein. Cell 181, 281–292, DOI: 10.1016/j.cell.2020.02.058 (2020).

10. Walls, A. C. et al. Unexpected Receptor Functional Mimicry Elucidates Activation of Coronavirus Fusion. Cell 176, 1026–1039, DOI: 10.1016/j.cell.2018.12.028 (2019).

11. Jin, Z. et al. Structure of Mpro from SARS-CoV-2 and discovery of its inhibitors. Nature 582, 289–293, DOI: 10.1038/s41586-020-2223-y (2020).

12. Anand, K., Ziebuhr, J., Wadhwani, P., Mesters, J. R. & Hilgenfeld, R. Coronavirus main proteinase (3CLpro) Structure: Basis for design of anti-SARS drugs. Science 300, 1763–1767, DOI: 10.1126/science.1085658 (2003).

13. Tan, J. et al. pH-dependent conformational flexibility of the SARS-CoV main proteinase (Mpro) dimer: Molecular dynamics simulations and multiple X-ray structure analyses. J. Mol. Biol. 354, 25–40, DOI: 10.1016/j.jmb.2005.09.012 (2005).

14. Xue, X. et al. Structures of Two Coronavirus Main Proteases: Implications for Substrate Binding and Antiviral Drug Design. J. Virol. 82, 2515–2527, DOI: 10.1128/jvi.02114-07 (2008).

15. Goyal, B. & Goyal, D. Targeting the Dimerization of the Main Protease of Coronaviruses: A Potential Broad-Spectrum Therapeutic Strategy. ACS combinatorial science 22, 297–305, DOI: 10.1021/acscombsci.0c00058 (2020).

16. Jin, Z. et al. Structure of M pro from COVID-19 virus and discovery of its inhibitors. Nature DOI: 10.1038/s41586-020-2223-y (2020).

17. Lee, Y., Lazim, R., Macalino, S. J. Y. & Choi, S. Importance of protein dynamics in the structure-based drug discovery of class A G protein-coupled receptors (GPCRs), DOI: 10.1016/j.sbi.2019.03.015 (2019).

18. Amaral, M. et al. Protein conformational flexibility modulates kinetics and thermodynamics of drug binding. Nat. Commun. 8, 1–14, DOI: 10.1038/s41467-017-02258-w (2017).

19. Jimenez-Roldan, J. E., Freedman, R. B., Römer, R. A. & Wells, S. A. Rapid simulation of protein motion: merging flexibility, rigidity and normal mode analyses. Phys. Biol. 9, 016008, DOI: 10.1088/1478-3975/9/1/016008 (2012).

20. Thorpe, M., Lei, M., Rader, A., Jacobs, D. J. & Kuhn, L. A. Protein flexibility and dynamics using constraint theory. J. Mol. Graph. Model. 19, 60–69, DOI: 10.1016/S1093-3263(00)00122-4 (2001).

21. Krebs, W. G. et al. Normal mode analysis of macromolecular motions in a database framework: Developing mode concentration as a useful classifying statistic. Proteins: Struct. Funct. Genet. 48, 682–695, DOI: 10.1002/prot.10168 (2002).

22. Tama, F. & Sanejouand, Y.-H. Conformational change of proteins arising from normal mode calculations. Protein Eng. Des. Sel. 14, 1–6, DOI: 10.1093/protein/14.1.1 (2001).

23. Suhre, K. & Sanejouand, Y.-H. On the potential of normal mode analysis for solving difficult molecular replacement problems. Acta Cryst D 60, 796–799 (2004).

24. Suhre, K. & Sanejouand, Y.-H. ElNemo: a normal mode web server for protein movement analysis and the generation of templates for molecular replacement. Nucleic Acids Res. 32, W610–W614 (2004).

25. Dykeman, E. C. & Sankey, O. F. Normal mode analysis and applications in biological physics. J. Physics: Condens. Matter 22, 423202, DOI: 10.1088/0953-8984/22/42/423202 (2010).

26. Wells, S., Menor, S., Hespenheide, B. & Thorpe, M. F. Constrained geometric simulation of diffusive motion in proteins. Phys. Biol. 2, S127–S136, DOI: 10.1088/1478-3975/2/4/S07 (2005).

27. Jolley, C. C., Wells, S. A., Hespenheide, B. M., Thorpe, M. F. & Fromme, P. Docking of Photosystem I Subunit C Using a Constrained Geometric Simulation. J. Am. Chem. Soc. 128, 8803–8812, DOI: 10.1021/ja0587749 (2006).

28. Li, H. et al. Protein flexibility is key to cisplatin crosslinking in calmodulin. Protein Sci. 21, 1269–1279, DOI: 10.1002/pro.2111 (2012).

29. Heal, J. W., Jimenez-Roldan, J. E., Wells, S. A., Freedman, R. B. & Römer, R. A. Inhibition of HIV-1 protease: the rigidity perspective. Bioinformatics 28, 350–357, DOI: 10.1093/bioinformatics/btr683 (2012).

30. Wells, S. A., van der Kamp, M. W., McGeagh, J. D. & Mulholland, A. J. Structure and Function in Homodimeric Enzymes: Simulations of Cooperative and Independent Functional Motions. PLOS ONE 10, e0133372, DOI: 10.1371/journal.pone.0133372 (2015).

31. Römer, R. A. et al. The flexibility and dynamics of protein disulfide isomerase. Proteins: Struct. Funct. Bioinforma. 84, 1776–1785, DOI: 10.1002/prot.25159 (2016).

32. Freedman, R. B. et al. ‘Something in the way she moves’: The functional significance of flexibility in the multiple roles of protein disulfide isomerase (PDI). Biochimica et Biophys. Acta (BBA) - Proteins Proteomics 1865, 1383–1394, DOI: 10.1016/j.bbapap.2017.08.014 (2017).

33. Farrell, D. W., Speranskiy, K. & Thorpe, M. F. Generating stereochemically acceptable protein pathways. Proteins: Struct. Funct. Bioinforma. 78, 2908–2921, DOI: 10.1002/prot.22810 (2010).

34. Ahmed, A. & Gohlke, H. Multiscale modeling of macromolecular conformational changes combining concepts from rigidity and elastic network theory. Proteins: Struct. Funct. Bioinforma. 63, 1038–1051, DOI: 10.1002/prot.20907 (2006).

35. Berman, H. M. et al. The Protein Data Bank. Nucleic Acids Res. 28, 235–242, DOI: 10.1093/nar/28.1.235 (2000).

36. Roemer, R. A., Roemer, N. S. & Wallis, A. K. Flex-Covid19 data repository, https://warwick.ac.uk/flex-covid19-data (2020).

37. Roemer, R. A. & Ratamero, E. M. “pdb2movie” github repository, https://github.com/RudoRoemer/pdb2movie (2020).

38. Roemer, R. A., Ratamero, E. M. & Moffat, S. PDB2Movie, https://pdb2movie.warwick.ac.uk (2020).

39. Word, J. M., Lovell, S. C., Richardson, J. S. & Richardson, D. C. Asparagine and glutamine: Using hydrogen atom contacts in the choice of side-chain amide orientation. J. Mol. Biol. 285, 1735–1747, DOI: 10.1006/jmbi.1998.2401 (1999).

40. DeLano, W. The PyMOL molecular graphics visualization programme.

41. Dahiyat, B. I., Benjamin Gordon, D. & Mayo, S. L. Automated design of the surface positions of protein helices. Protein Sci. 6, 1333–1337, DOI: 10.1002/pro.5560060622 (1997).

42. Wells, S. A., Jimenez-Roldan, J. E. & Römer, R. A. Comparative analysis of rigidity across protein families. Phys. Biol. 6, 046005, DOI: 10.1088/1478-3975/6/4/046005 (2009).

43. Tirion, M. M. Large Amplitude Elastic Motions in Proteins from a Single-Parameter, Atomic Analysis. Phys. Rev. Lett. 77, 1905–1908, DOI: 10.1103/PhysRevLett.77.1905 (1996).

