## supplementary materials for "Flexibility and mobility of SARS-CoV-2-related protein structures"

### *Supplemental Material for*
### Flexibility and Mobility of SARS-CoV-2-related protein structures

**Rudolf A. Römer**[1,2,3*], **Navodya S. Römer**[4], **and A. Katrine Wallis**[5]

[1]CY Advanced Studies and LPTM (UMR8089 of CNRS), CY Cergy-Paris Université, F-95302 Cergy-Pontoise, France
[2]Department of Physics, University of Warwick, Coventry, CV4 7AL, United Kingdom
[4]School of Life Sciences, University of Lincoln, Brayford Pool Campus, Lincoln, LN6 7TS, United Kingdom
[5]School of Life Sciences, University of Warwick, Coventry, CV4 7AL, United Kingdom
[*]

#### ABSTRACT

#### 1 Sources and video links

In Table S1 we give the complete list of all PDB codes used in our study, link their supporting publications as well as the list of modes and $E_{\mathrm{cut}}$ values used in computing the full motion movies available at Ref. 1. We also provide the full motion movies for the structures given in the main text in Fig. 1, namely the monomer of the SARS-Cov-2 spike ecto domain (6vxx)[?] and the trimer of main protease[3]. Similarly, for main text Fig. 2 we include the movies of the closed SARS-Cov-2 spike ecto domain trimer in side[4] and top view.[5] Last, for the open SARS-Cov-2 spike ecto domain trimer in main text Fig. 3, we again give the side[6] and top views.[7]

| PDB code | Ref. | rigidity class. | $E_{cut}$'s (kcal/mol) | protein type | role | oligomeric state | error |
|---|---|---|---|---|---|---|---|
| The following PDB files were downloaded 18 April 2020 | | | | | | | |
| 5r7y | 8 | 1st | 1, 2, 3, 0.2 | Main protease | cleavage of poly protein | dimer | |
| 5r7z | 8 | 1st | 1, 2, 3 | Main protease | icw. Z1220452176 | dimer | |
| 5r80 | 8 | 1st | 1, 2, 3 | Main protease | icw. Z18197050 | dimer | |
| 5r81 | 8 | 2nd | 1, 2, 3 | Main protease | icw. Z1367324110 | dimer | |
| 5r82 | 8 | 2nd | 1, 2, 3 | Main protease | icw. Z219104216 | dimer | |
| 5r83 | 8 | 1st | 1, 2, 3 | Main protease | icw. Z44592329 | dimer | |
| 5r84 | 8 | 1st | 1, 2, 3, 2.5 | Main protease | icw. Z31792168 | dimer | |
| 5r8t | 8 | 1st | 1, 2, 3 | Main protease | | dimer | |
| 5re4 | 8 | 1st | 1, 2, 3, 1.1 | Main protease | icw. Z1129283193 | dimer | |
| 5re5 | 8 | 1st | 1, 2, 3 | Main protease | icw. Z1129283193 | dimer | |
| 5re6 | 8 | 1st | 1, 2, 3 | Main protease | icw. Z1129283193 | dimer | |
| 5re7 | 8 | | 1, 2, 3 | Main protease | icw. Z30932204 | dimer | bond distance between intraresidue atoms 945 and 950 exceeds 6 Å |
| 5re8 | 8 | 1st | 1, 2, 3 | Main protease | icw. Z2737076969 | dimer | |
| 5re9 | 8 | | 1, 2, 3 | Main protease | icw. Z2856434836 | dimer | bond distance between intraresidue atoms 945 and 950 exceeds 6 Å |
| 5rea | 8 | | 1, 2, 3 | Main protease | icw. Z31432226 | dimer | bond distance between intraresidue atoms 954 and 979 exceeds 6 Å |
| 5reb | 8 | 1st | 1, 2, 3 | Main protease | icw. Z2856434899 | dimer | |
| 5rec | 8 | 1st | 1, 2, 3, 0.4 | Main protease | icw. Z1587220559 | dimer | |
| 5red | 8 | | 1, 2, 3 | Main protease | icw. Z2856434865 | dimer | bond distance between intraresidue atoms 952 and 957 exceeds 6 Å |
| 5ree | 8 | 1st | 1, 2, 3 | Main protease | icw. Z2217052426 | dimer | |
| 5ref | 8 | 1st | 1, 2, 3 | Main protease | icw. Z24758179 | dimer | |
| 5reg | 8 | | 1, 2, 3 | Main protease | icw. Z1545313172 | dimer | bond distance between intraresidue atoms 947 and 972 exceeds 6 Å |
| 5reh | 8 | 1st | 1, 2, 3, 1.8 | Main protease | icw. Z111507846 | dimer | |
| 5rei | 8 | 1st | 1, 2, 3 | Main protease | icw. Z2856434856 | dimer | |
| 5rej | 8 | 1st | 1, 2, 3 | Main protease | icw. PCM-0102241 | dimer | |
| 5rek | 8 | 1st | 1, 2, 3 | Main protease | icw. PCM-0102327 | dimer | |

continued on next page

| PDB code | Ref. | rigidity class. | $E_{cut}$'s (kcal/mol) | protein type | role | oligomeric state | error |
|---|---|---|---|---|---|---|---|
| 5rel | 8 | 2nd | 1, 2, 3, 1.5 | Main protease | icw. PCM-0102340 | dimer | |
| 5rem | 8 | 1st | 1, 2, 3 | Main protease | icw. PCM-0103016 | dimer | |
| 5ren | 8 | 1st | 1, 2, 3 | Main protease | icw. PCM-0102425 | dimer | |
| 5reo | 8 | 1st | 1, 2, 3, 1.7 | Main protease | icw. PCM-0102578 | dimer | |
| 5rep | 8 | 1st | 1, 2, 3 | Main protease | icw. PCM-0102201 | dimer | |
| 5rer | 8 | 1st | 1, 2, 3 | Main protease | icw. PCM-0102615 | dimer | |
| 5res | 8 | 1st | 1, 2, 3 | Main protease | icw. PCM-0102281 | dimer | |
| 5ret | 8 | 1st | 1, 2, 3 | Main protease | icw. PCM-0102269 | dimer | |
| 5reu | 8 | 1st | 1, 2, 3 | Main protease | icw. PCM-0102395 | dimer | |
| 5rev | 8 | 1st | 1, 2, 3 | Main protease | icw. PCM-0103072 | dimer | |
| 5rew | 8 | 1st | 1, 2, 3 | Main protease | icw. PCM-0102275 | dimer | |
| 5rex | 8 | 1st | 1, 2, 3 | Main protease | icw. PCM-0102287 | dimer | |
| 5rey | 8 | 1st | 1, 2, 3, 0.5 | Main protease | icw. PCM-0102911 | dimer | |
| 5rez | 8 | 1st | 1, 2, 3 | Main protease | icw. POB0129 | dimer | |
| 5rf0 | 8 | 1st | 1, 2, 3 | Main protease | icw. POB0073 | dimer | |
| 5rf1 | 8 | 1st | 1, 2, 3 | Main protease | icw. NCL-00023830 | dimer | |
| 5rf2 | 8 | 1st | 1, 2, 3 | Main protease | icw. Z1741969146 | dimer | |
| 5rf3 | 8 | 1st | 1, 2, 3 | Main protease | icw. Z1741970824 | dimer | |
| 5rf4 | 8 | 1st | 1, 2, 3 | Main protease | icw. Z1741982125 | dimer | |
| 5rf5 | 8 | 1st | 1, 2, 3, 1.5 | Main protease | icw. Z3241250482 | dimer | |
| 5rf6 | 8 | 2nd | 1, 2, 3 | Main protease | icw. Z1348371854 | dimer | |
| 5rf7 | 8 | 1st | 1, 2, 3 | Main protease | icw. Z316425948_minor | dimer | |
| 5rf8 | 8 | 1st | 1, 2, 3 | Main protease | icw. Z271004858 | dimer | |
| 5rf9 | 8 | 1st | 1, 2, 3 | Main protease | icw. Z217038356 | dimer | |
| 5rfa | 8 | 1st | 1, 2, 3 | Main protease | icw. Z2643472210 | dimer | |
| 5rfb | 8 | 1st | 1, 2, 3 | Main protease | icw. Z1271660837 | dimer | |
| 5rfc | 8 | 1st | 1, 2, 3 | Main protease | icw. Z979145504 | dimer | |
| 5rfd | 8 | 1st | 1, 2, 3 | Main protease | icw. Z126932614 | dimer | |
| 5rfe | 8 | 1st | 1, 2, 3 | Main protease | icw. Z126932614 | dimer | |
| 5rff | 8 | 1st | 1, 2, 3 | Main protease | icw. PCM-0102704 | dimer | |
| 5rfg | 8 | 1st | 1, 2, 3 | Main protease | icw. PCM-0102372 | dimer | |
| 5rfh | 8 | 1st | 1, 2, 3 | Main protease | icw. PCM-0102277 | dimer | |
| 5rfi | 8 | 1st | 1, 2, 3, 1.5 | Main protease | icw. PCM-0102353 | dimer | |
| 5rfj | 8 | 1st | 1, 2, 3, 1.7 | Main protease | icw. PCM-0103067 | dimer | |
| 5rfk | 8 | 1st | 1, 2, 3 | Main protease | icw. PCM-0102575 | dimer | |
| 5rfl | 8 | 1st | 1, 2, 3 | Main protease | icw. PCM-0102389 | dimer | |

continued on next page

| PDB code | Ref. | rigidity class. | $E_{cut}$'s (kcal/mol) | protein type | role | oligomeric state | error |
|---|---|---|---|---|---|---|---|
| 5rfm | 8 | 1st | 1, 2, 3, 0.5 | Main protease | icw. PCM-0102539 | dimer | |
| 5rfn | 8 | 1st | 1, 2, 3, 1.9 | Main protease | icw. PCM-0102868 | dimer | |
| 5rfo | 8 | 1st | 1, 2, 3 | Main protease | n complex with PCM-0102972 | dimer | |
| 5rfp | 8 | 2nd | 1, 2, 3 | Main protease | icw. PCM-0102190 | dimer | |
| 5rfq | 8 | 1st | 1, 2, 3, 1.8 | Main protease | icw. PCM-0102179 | dimer | |
| 5rfr | 8 | 1st | 1, 2, 3 | Main protease | icw. PCM-0102169 | dimer | |
| 5rfs | 8 | 1st | 1, 2, 3 | Main protease | icw. PCM-0102739 | dimer | |
| 5rft | 8 | 1st | 1, 2, 3 | Main protease | icw. PCM-0102432 | dimer | |
| 5rfu | 8 | 1st | 1, 2, 3 | Main protease | icw. PCM-0102121 | dimer | |
| 5rfv | 8 | 1st | 1, 2, 3 | Main protease | icw. PCM-0102306 | dimer | |
| 5rfw | 8 | 1st | 1, 2, 3, 1.5 | Main protease | icw. PCM-0102243 | dimer | |
| 5rfx | 8 | 1st | 1, 2, 3 | Main protease | icw. PCM-0102254 | dimer | |
| 5rfy | 8 | 1st | 1, 2, 3, 0.5 1.5 | Main protease | icw. PCM-0102974 | dimer | |
| 5rfz | 8 | 1st | 1, 2, 3 | Main protease | icw. PCM-0102274 | dimer | |
| 5rg0 | 8 | 1st | 1, 2, 3 | Main protease | icw. PCM-0102535 | dimer | |
| 5rg1 | 8 | 1st | 1, 2, 3, 1.5 | Main protease | icw. NCL-00024905 | dimer | |
| 5rg2 | 8 | 2nd | 1, 2, 3, 2.5 | Main protease | icw. NCL-00025058 | dimer | |
| 5rg3 | 8 | 1st | 1, 2, 3 | Main protease | icw. NCL-00025412 | dimer | |
| 5rgg | 8 | 1st | 1, 2, 3, 1.5 | Main protease | icw. Z2856434890 (Mpro-x0165) | dimer | |
| 5rgh | 8 | 1st | 1, 2, 3 | Main protease | icw. Z1619978933 (Mpro-x0395) | dimer | |
| 5rgi | 8 | 1st | 1, 2, 3 | Main protease | icw. Z369936976 (Mpro-x0397) | dimer | |
| 5rgj | 8 | 2nd | 1, 2, 3, 1.5 | Main protease | icw. Z1401276297 (Mpro-x0425) | dimer | |
| 5rgk | 8 | 1st | 1, 2, 3 | Main protease | icw. Z1310876699 (Mpro-x0426 | dimer | |
| 5rgl | 8 | 1st | 1, 2, 3, 1.6 | Main protease | icw. PCM-0102962 (Mpro-x0705) | dimer | |
| 5rgm | 8 | 1st | 1, 2, 3 | Main protease | icw. PCM-0102142 (Mpro-x0708) | dimer | |
| 5rgn | 8 | 1st | 1, 2, 3, 1.5 | Main protease | icw. PCM-0102759 (Mpro-x0731) | dimer | |

continued on next page

| PDB code | Ref. | rigidity class. | $E_{\mathrm{cut}}$'s (kcal/mol) | protein type | role | oligomeric state | error |
|---|---|---|---|---|---|---|---|
| 5rgo | 8 | 1st | 1, 2, 3 | Main protease | icw. PCM-0102248 (Mpro-x0736) | dimer | |
| 5rgp | 8 | 1st | 1, 2, 3 | Main protease | icw. PCM-0102628 (Mpro-x0771) | dimer | |
| 5rgq | 8 | 1st | 1, 2, 3, 1.75 | Main protease | icw. Z1849009686 (Mpro-x1086) | dimer | |
| 5rgr | 8 | 1st | 1, 2, 3 | Main protease | icw. Z328695024 (Mpro-x1101) | dimer | |
| 5rgs | 8 | 1st | 1, 2, 3 | Main protease | icw. Z1259086950 (Mpro-x1163) | dimer | |
| 6lu7 | 9 | 1st | 1, 2, 3, 0.7 1.5 | Main protease | icw. an inhibitor N3 | dimer | |
| 6lvn | 10 | brick | 1, 2, 3 | Spike protein HR2 domain | Spike protein does binding to receptor and involved in fusion of membranes | tetramer | |
| 6lxt | 11 | 1st | 1, 2, 3, 3.3 3.6 4.6 | Spike protein post fusion core | S2 subunit | trimer | |
| 6lzg | 12 | 1st | 1, 2, 3 | Spike protein domain icw. ace2 | | dimer | |
| 6m03 | 13 | 1st | 1, 2, 3 | main protease | apoform | dimer | |
| 6m0j | 14 | | 1, 2, 3 | Spike protein domain icw. ace2 | | hetero-dimer | bond distance between intraresidue atoms 3327 and 3342 exceeds 6 Å |
| 6m17 | 15 | domain | 1, 2, 3, 0.1 0.2 0.4 0.5 0.7 | Ace2 icw. amino acid transporte and spike protein domain | | hetero-hexamer | |
| 6m18 | 15 | domain | 1, 2, 3, 0.03 0.06 0.2 0.5 | ace2 icw. amino acid transporter | This structure contains no viral protein | hetero-tetramer | |
| 6m1d | 15 | 2nd | 1, 2, 3, 0.2 0.4 0.6 | ace2 icw. amino acid transporter | This structure contains no viral protein | hetero-tetramer | |
| 6m2n | 16 | 2nd | 1, 2, 3, 0.5 | Protease | structure with ligand | dimer | |
| 6m2q | 17 | 2nd | 1, 2, 3 | Protease | apo protein | dimer | |
| 6m3m | 18 | 2nd | 1, 2, 3, 0.15 | Nucleocapside protein RNA binding domain | | monomer | |

continued on next page

| PDB code | Ref. | rigidity class. | $E_{cut}$'s (kcal/mol) | protein type | role | oligomeric state | error |
|---|---|---|---|---|---|---|---|
| 6m71 | 19 | 2nd | 1, 2, 3, 0.08 0.17 0.4 0.7 | RNA polymerase | icw. cofactors | hetero-tetramer | |
| 6vsb | 20 | domain | 1, 2, 3, 0.005 0.04 0.1 0.35 0.8 1.5 | Spike ecto domain | EM structure 3.5 angstrom | trimer | |
| 6vw1 | 21 | 1st | 1, 2, 3, 0.5 4.0 4.5 | Spike receptor binding domain with Ace2 | | hetero-dimer | |
| 6vww | 22 | | 1, 2, 3 | NSP15 endoribonuclease | | hexamer | bond distance between intraresidue atoms 26 and 27 exceeds 6 Å |
| 6vxs | 23 | domain | 1, 2, 3, 1.3 | ADP ribose phosphotase | | monomer | |
| 6vxx | 24 | domain | 1, 2, 3, 0.001 0.08 0.5 1.5 | Spike ecto domain | EM structure 2.8 angstrom closed | trimer | |
| 6vyb | 24 | domain | 1, 2, 3, 0.001 0.5 | Spike ecto domain | Em structure open structure 3.2 angstrom | trimer | |
| 6vyo | 23 | brick | 1, 2, 3, 0.1 0.7 | nucleocapside phosphoprotein | | tetramer | |
| 6w01 | 22 | | 1, 2, 3 | NSP15 endoribonuclease | in the Complex with a Citrate | hexamer | bond distance between intraresidue atoms 9 and 24 exceeds 6 Å |
| 6w02 | 23 | | 1, 2, 3 | ADP ribose phosphotase | complex with ADP ribose | monomer | bond distance between intraresidue atoms 2095 and 2100 exceeds 6 Å |
| 6w41 | 25 | | 1, 2, 3 | Spike receptor binding domain complexed with antibody | icw. human antibody CR3022 | hetero-trimer | steric clashes |
| 6w4b | 23 | 1st | 1, 2, 3, 0.1 0.4 | NSP9 RNA binding protein | | dimer | |
| 6w4h | 23 | | 1, 2, 3 | NSP16 - NSP10 Complex | | hetero-dimer | bond distance between intraresidue atoms 430 and 453 exceeds 6 Å |

continued on next page

| PDB code | Ref. | rigidity class. | $E_{cut}$'s (kcal/mol) | protein type | role | oligomeric state | error |
|---|---|---|---|---|---|---|---|
| 6w61 | 23 | | 1, 2, 3 | NSP10 and NSP16 complex: methyltransferase stimulatory complex | | hetero-dimer | bond distance between intraresidue atoms 320 and 345 exceeds 6 Å |
| 6w63 | 23 | 2nd | 1, 2, 3, 1.5 | Main protease | bound to potent broad-spectrum non-covalent inhibitor X77 | dimer | |
| 6w6y | 23 | 2nd | 1, 2, 3 | ADP ribose phosphotase | icw. AMP | monomer | |
| 6w75 | 23 | | 1, 2, 3 | NSP10-NSP16 complex | | hetero-dimer | bond distance between intraresidue atoms 322 and 323 exceeds 6 Å |
| 6w9c | 23 | 1st | 1, 2, 3, 0.5 | papain like protease | | trimer | |
| 6w9q | 26 | 1st | 1, 2, 3 | NSP9 RNA replicase | | dimer | |
| 6wcf | 23 | | 1, 2, 3 | ADP ribose phosphotase | icw. MES | monomer | bond distance between intraresidue atoms 1189 and 1214 exceeds 6 Å |
| 6wen | 23 | | 1, 2, 3 | ADP ribose phosphotase | apo form | monomer | bond distance between intraresidue atoms 1141 and 1147 exceeds 6 Å |
| 6y2e | 13 | 2nd | 1, 2, 3 | Main protease | | dimer | |
| 6y2f | 13 | 1st | 1, 2, 3 | Main protease | monoclinic form | dimer | |
| 6y2g | 13 | 2nd | 1, 2, 3, 0.5 1.5 | Main protease | orthorhombic form | dimer | |
| 6y84 | 27 | | 1, 2, 3 | Main protease | unliganded active site | dimer | bond distance between intraresidue atoms 4600 and 4603 exceeds 6 Å |
| 6yb7 | 27 | 2nd | 1, 2, 3, 2.5 4.0 | Main protease | supercedes 6y84 | dimer | |
| 6yi3 | 28 | | 1, 2, 3 | nucleocapside phosphoprotein | NMR structure | | bond distance between intraresidue atoms 4600 and 4603 exceeds 6 Å |

continued on next page

| PDB code | Ref. | rigidity class. | $E_{cut}$'s (kcal/mol) | protein type | role | oligomeric state | error |
|---|---|---|---|---|---|---|---|
| 6yla | 29 | 2nd | 1, 2, 3, 0.7 1.5 | spike receptor binding domain with Fab fragment | | hetero-trimer | |
| 7btf | 19 | 2nd | 1, 2, 3, 0.1 0.3 0.4 0.8 | RNA polymerase | icw. cofactors in reduced condition | hetero-tetramer | |
| The following PDB files were downloaded 19 May 2020 | | | | | | | |
| 3r24 | 18 | 1st | 1, 2, 3 | transferase | RNA maturation | hetero-dimer | |
| 6lze | 13 | 2nd | 1, 2, 3, 1.7 | | icw. an inhibitor 11a | dimer | |
| 6m0k | 13 | 2nd | 1, 2, 3 | main protease | icw. an inhibitor 11b | dimer | |
| 6w37 | 23 | brick | 1, 2, 3 | | | monomer | |
| 6w7y | 18 | domain | 1, 2, 3, 0.015 0.15 0.4 0.6 | | SARS-CoV-2 reactive human antibody CR3022 | dimer | |
| 6wey | 30 | 1, 2, 3, 2.5 4.0 5.0 | | | | monomer | |
| 6wiq | 23 | "3,2" | 1, 2, 3, 0.001 0.03 0.5 1.5 | | | hetero-tetramer | |
| 6wji | 23 | domain | 1, 2, 3, 0.001 0.13 0.6 1.7 | | Crystal Structure of C-terminal Dimerization Domain of Nucleocapsid Phosphoprotein from SARS-CoV-2 | dimer | |
| 6wjt | 23 | | 1, 2, 3 | | | hetero-dimer | bond distance between intraresidue atoms 01952 and 01967 exceeds 6 Å |
| 6wkp | 23 | brick | 1, 2, 3, 0.08 0.5 | | monoclinic crystal form | tetramer | |
| 6wkq | 23 | 2nd | 1, 2, 3 | | icw. Sinefungin | hetero-dimer | |
| 6wks-bundle1 | 31 | 2nd | 1, 2, 3, 0.01 0.2 0.5 1.5 | | | hetero-tetramer | |

continued on next page

| PDB code | Ref. | rigidity class. | $E_{cut}$'s (kcal/mol) | protein type | role | oligomeric state | error |
|---|---|---|---|---|---|---|---|
| 6wlc | 23 | | 1, 2, 3 | | in the Complex with Uridine-5'-Monophosphate | hexamer | bond distance between intraresidue atoms 1 and 4 exceeds 6 Å |
| 6wnp | 23 | | 1, 2, 3 | | bound to Boceprevir at 1.45 A | dimer | bond distance between intraresidue atoms 04378 and 04381 exceeds 6 Å |
| 6woj | 23 | 2nd | 1, 2, 3, 4 | | icw. ADP-ribose | monomer | |
| 6wq3 | 23 | | 1, 2, 3 | | icw. 7-methyl-GpppA and S-adenosyl-L-homocysteine. | hetero-dimer | bond distance between intraresidue atoms 03495 and 03516 exceeds 6 Å |
| 6wqd | 23 | | 1, 2, 3 | | Complex of NSP7 and the C-terminal Domain of NSP8 from SARS-CoV-2 | hetero-tetramer | bond distance between intraresidue atoms 359 and 376 exceeds 6 Å |
| 6wqf | 32 | | 1, 2, 3 | | Revealed by Room Temperature X-ray Crystallography | dimer | bond distance between intraresidue atoms 03306 and 03331 exceeds 6 Å |
| 6wrh | 23 | | 1, 2, 3 | | | monomer | bond distance between intraresidue atoms 02089 and 02110 exceeds 6 Å |
| 6wrz | 23 | | 1, 2, 3 | | | hetero-dimer | bond distance between intraresidue atoms 03491 and 03510 exceeds 6 Å |
| 6wtc | 23 | domain | 1, 2, 3, 4 | | Second Form of the Co-factor Complex of NSP7 and the C-terminal Domain of NSP8 from SARS CoV-2 | hetero-tetramer | |
| 6wvn | 23 | 2nd | 1, 2, 3 | | icw. 7-methyl-GpppA and S-Adenosylmethionine. | hetero-dimer | |

continued on next page

| PDB code | Ref. | rigidity class. | $E_{cut}$'s (kcal/mol) | protein type | role | oligomeric state | error |
|---|---|---|---|---|---|---|---|
| 6y7m-bundle1 | 13 | 2nd | 1, 2, 3, 0.3 0.4 | | | dimer | |
| 6yhu | 33 | domain | 1, 2, 3 | nsp7-nsp8 complex of SARS-CoV-2 | | hetero-dimer | |
| 6ym0 | 34 | | 1, 2, 3 | | icw. CR3022 Fab | hetero-trimer | steric clashes |
| 6ynq | 35 | | 1, 2, 3 | main protease | bound to 2-Methyl-1-tetralone | dimer | bond distance between intraresidue atoms 062 and 085 exceeds 6 Å |
| 6yor | 36 | 2nd | 1, 2, 3, 0.26 | SARS-CoV-2 spike S1 protein | icw. CR3022 Fab | hetero-trimer | |
| 6yt8 | 37 | | 1, 2, 3 | main protease | bound to pyrithione zinc | dimer | bond distance between intraresidue atoms 079 and 082 exceeds 6 Å |
| 6yva | 38 | 1st | 1, 2, 3, 0.15 0.3 1.5 | | | hetero-dimer | |
| 6ywk | 39 | domain | 1, 2, 3, 1.5 | SARS-CoV-2 (Covid-19) NSP3 macrodomain | icw. HEPES | monomer | |
| 6ywl | 39 | domain | 1, 2, 3, 1.5 | SARS-CoV-2 (Covid-19) NSP3 macrodomain | icw. ADP-ribose | monomer | |
| 6ywm | 39 | domain | 1, 2, 3, 2.5 4.0 | SARS-CoV-2 (Covid-19) NSP3 macrodomain | icw. MES | monomer | |
| 6yyt | 40 | 2nd | 1, 2, 3, 0.004 0.15 0.3 0.4 | replicating SARS-CoV-2 polymerase | | hetero-tetramer | |
| 6yz1 | 41 | | 1, 2, 3 | | complex with Sinefungin | hetero-dimer | bond distance between intraresidue atoms 03770 and 03773 exceeds 6 Å |
| 7bqy | 9 | 2nd | 1, 2, 3, 0.5 1.2 | main protease | icw. AN INHIBITOR N3 at 1.7 angstrom | dimer | |
| 7bro | 42 | 2nd | 1, 2, 3, 1.3 1.5 | main protease | | dimer | |

continued on next page

| PDB code | Ref. | rigidity class. | $E_{cut}$'s (kcal/mol) | protein type | role | oligomeric state | error |
|---|---|---|---|---|---|---|---|
| 7brp | 42 | 2nd | 1, 2, 3, 0.5 0.75 1.3 | main protease | complexed with Boceprevir | dimer | |
| 7brr | 42 | 1st | 1, 2, 3, 1.5 | main protease | complexed with GC376 | dimer | |
| 7buy | 9 | 2nd | 1, 2, 3, 0.5 | main protease | icw. carmofur | dimer | |
| 7bv1 | 43 | 2nd | 1, 2, 3, 0.05 0.15 0.25 0.5 1.5 | nsp12-nsp7-nsp8 complex | apo? | hetero-tetramer | |
| 7bv2 | 43 | 2nd | 1, 2, 3, 0.04 0.15 0.4 | nsp12-nsp7-nsp8 complex | bound to the template-primer RNA and triphosphate form of Remdesivir(RTP) | hetero-trimer | |
| 7bz5 | 44 | brick | 1, 2, 3 | COVID-19 virus spike receptor-binding domain | complexed with a neutralizing antibody | hetero-trimer | |
| The following PDB files were downloaded 29 May 2020 | | | | | | | |
| 5rgt | 8 | | 1, 2, 3 | main protease | icw. Z4439011607 | dimer | The bond distance between intraresidue atoms 03302 and 03325 exceeds 6 Å |
| 5rgu | 8 | 1st | 1, 2, 3, 2.5 | main protease | icw. Z4444622180 | dimer | |
| 5rgv | 8 | 1st | 1, 2, 3 | main protease | icw. Z4444622066 | dimer | |
| 5rgw | 8 | 1st | 1, 2, 3 | main protease | icw. Z4444621910 | dimer | |
| 5rgx | 8 | 1st | 1, 2, 3 | main protease | icw. Z1344037997 | dimer | |
| 5rgy | 8 | brick | 1, 2, 3, 1.7 1.9 | main protease | icw. Z1535580916 | dimer | |
| 5rgz | 8 | 1st | 1, 2, 3 | main protease | icw. Z1343543528 | dimer | |
| 5rh0 | 8 | 1st | 1, 2, 3 | main protease | icw. Z1286870272 | dimer | |
| 5rh1 | 8 | 1st | 1, 2, 3, 1.7 | main protease | icw. Z2010253653 | dimer | |
| 5rh2 | 8 | 2nd | 1, 2, 3 | main protease | icw. Z1129289650 | dimer | |
| 5rh3 | 8 | | 1, 2, 3 | main protease | icw. Z1264525706 | dimer | The bond distance between intraresidue atoms 03302 and 03325 exceeds 6 Å |

continued on next page

| PDB code | Ref. | rigidity class. | $E_{cut}$'s (kcal/mol) | protein type | role | oligomeric state | error |
|---|---|---|---|---|---|---|---|
| 5rh4 | 8 | | 1, 2, 3 | main protease | icw. Z1530425063 | dimer | The bond distance between intraresidue atoms 02536 and 02551 exceeds 6 Å |
| 5rh5 | 8 | 1st | 1, 2, 3, 2.1 2.4 | main protease | icw. Z4439011520 | dimer | |
| 5rh6 | 8 | 2nd | 1, 2, 3 | main protease | icw. Z4439011588 | dimer | |
| 5rh7 | 8 | | 1, 2, 3 | main protease | icw. Z4439011584 | dimer | The bond distance between intraresidue atoms 03310 and 03333 exceeds 6 Å |
| 5rh8 | 8 | 1st | 1, 2, 3, 1.5 | main protease | icw. Z4444621965 | dimer | |
| 5rh9 | 8 | | 1, 2, 3 | main protease | icw. Z4438424255 | dimer | The bond distance between intraresidue atoms 03302 and 03325 exceeds 6 Å |
| 5rha | 8 | | 1, 2, 3 | main protease | icw. Z147647874 | dimer | The bond distance between intraresidue atoms 03302 and 03327 exceeds 6 Å |
| 6wps | 45 | domain | 1, 2, 3 | SARS-CoV-2 spike glycoprotein | icw. the S309 neutralizing antibody Fab fragment | Hetero-9-mer | |
| 6wpt | 45 | domain | 1, 2, 3 | SARS-CoV-2 spike glycoprotein | icw. the S309 neutralizing antibody Fab fragment (open state) | Hetero-7-mer | |
| 6wtj | 46 | 2nd | 1, 2, 3 | main protease | Feline coronavirus drug inhibits | dimer | |
| 6wtk | 46 | 1st | 1, 2, 3 | main protease | Feline coronavirus drug inhibits | dimer | |
| 6wtm | 46 | 2nd | 1, 2, 3 | main protease | Feline coronavirus drug inhibits | dimer | |
| 6wtt | 47 | 2nd | 1, 2, 3 | main protease | with inhibitor GC-376 | dimer | |
| 6wuu | 48 | 1st | 1, 2, 3, 0.5 | Papain-like protease | icw. peptide inhibitor VIR250 | hetero-dimer | |
| 6wx4 | 48 | | 1, 2, 3 | Papain-like protease | icw. peptide inhibitor VIR251 | monomer | steric clashes |

continued on next page

| PDB code | Ref. | rigidity class. | $E_{cut}$'s (kcal/mol) | protein type | role | oligomeric state | error |
|---|---|---|---|---|---|---|---|
| 6wxc | 23 | | 1, 2, 3 | NSP15 Endoribonuclease | n the Complex with potential repurposing drug Tipiracil | hexamer | The bond distance between intraresidue atoms 1 and 4 exceeds 6 Å |
| 6wxd | 49 | brick | 1, 2, 3 | Nsp9 RNA-replicase | | dimer | |
| 6wzo | 50 | domain | 1, 2, 3, 0.5 | "Nucleocapsid dimerization domain, P1 form" | | dimer | |
| 6wzq | 50 | domain | 1, 2, 3 | "Nucleocapsid dimerization domain, P21 form" | | dimer | |
| 6wzu | 23 | | 1, 2, 3 | Papain-Like Protease | P3221 space group | monomer | The bond distance between intraresidue atoms 02023 and 02026 exceeds 6 Å |
| 6x1b | 23 | | 1, 2, 3 | NSP15 Endoribonuclease | in the Complex with the Product Nucleotide GpU. | hexamer | The bond distance between intraresidue atoms 05377 and 05380 exceeds 6 Å |
| 6x29 | 51 | domain | 1, 2, 3, 0.002 0.05 0.3 | | | trimer | |
| 6x2a | 51 | domain | 1, 2, 3, 0.005 0.1 | | | trimer | |
| 6x2b | 51 | domain | 1, 2, 3, 0.005 0.14 0.5 | | | trimer | |
| 6x2c | 51 | domain | 1, 2, 3, 0.001 0.05 0.25 0.7 | | | trimer | |
| 6yun | 52 | domain | 1, 2, 3 | C-terminal Dimerization Domain of Nucleocapsid Phosphoprotein | | dimer | The bond distance between intraresidue atoms 03404 and 03429 exceeds 6 Å |
| 6yvf | 35 | | 1, 2, 3 | Main Protease | bound to AZD6482 | dimer | steric clashes |

continued on next page

| PDB code | Ref. | rigidity class. | $E_{cut}$'s (kcal/mol) | protein type | role | oligomeric state | error |
|---|---|---|---|---|---|---|---|
| 6yz6 | 35 | 1st | 1, 2, 3 | | hemiacetal complex of Main Protease and Leupeptin | dimer | |
| 7bw4 | 53 | 2nd | 1, 2, 3, 0.015 0.15 | RNA-dependent RNA polymerase | | hetero-tetramer | |
| 7c01 | 54 | 2nd | 1, 2, 3 | | potent human neutralizing antibody targeting SARS-CoV-2 RBD | hetero-trimer | |
| 7c22 | 55 | domain | 1, 2, 3, 0.5 | C-terminal domain of SARS-CoV-2 nucleocapsid protein | | dimer | |
| 7c2i | 56 | 2nd | 1, 2, 3 | nsp16-nsp10 heterodimer | icw. SAM (with additional SAM during crystallization) | hetero-dimer | |
| 7c2j | 56 | 2nd | 1, 2, 3, 1.3 | nsp16-nsp10 heterodimer | icw. SAM (with additional SAM during crystallization) | hetero-dimer | |
| wuhan_ace2_MD | ? | domain | 1, 2, 3, 0.001 0.1 0.2 0.3 0.4 0.5 0.8 1.5 | | | | |
| wuhan_ace2_min | ? | domain | 1, 2, 3, 0.001 0.1 0.2 0.3 0.4 0.5 0.8 1.5 | | | | |
| wuhan_E_protein_3x29 | ? | "3,1" | 1, 2, 3, 0.2 0.5 | | | | |
| wuhan_spike (6vsb) | ? | "3,1" | 1, 2, 3, 0.001 0.1 0.2 0.3 0.4 0.5 0.8 1.5 | | | | |

**Table S1.** List of all SARS-Cov-2-related structures investigated in this study. The relevant references have been provided if they had been listed on the PDB download pages for each PDB code at the time of download access. The abbreviation "icw." stands for "in complex with". Two large scale collaborations have been labelled by acronyms PANDDA[8] and CSGID[23]. The rigidity association into solid "brick", 1st and 2nd order as well as rigidity domains are given on column 3. Values for $E_{cut}$ are given explicitly whereas all results have been computed for all 6 modes $m_7$ to $m_{12}$. Protein description, role and oligomeric state are as written of from the information in each PDB entry or in the referenced publication. The last column indicates the computational error encountered when now motion result has been computed.

#### References


1. Roemer, R. A., Roemer, N. S. & Wallis, A. K. Flex-Covid19 data repository https://warwick.ac.uk/flex-covid19-data (2020).
2. Roemer, R. A., Roemer, N. S. & Wallis, A. K. https://livewarwickac.sharepoint.com/:v:/s/Flex-Covid19/EQern4arlFtFmBJgcdJGZfoBRXlS4foHGWeL_CY-R8qDVA?e=JdhKHS (2020).
3. Roemer, R. A., Roemer, N. S. & Wallis, A. K. https://livewarwickac.sharepoint.com/:v:/s/Flex-Covid19/EW5Tj2CCMfJMop2RCxJNSFYBBbiFaeNLc9MC1eNfxX-lug?e=w5883E (2020).
4. Roemer, R. A., Roemer, N. S. & Wallis, A. K. https://livewarwickac.sharepoint.com/:v:/s/Flex-Covid19/EVRO00oUOFJGlnys98iQ3W0BeQPnxIH92GkwP9w_GuKzzg?e=XTLwT7 (2020).
5. Roemer, R. A., Roemer, N. S. & Wallis, A. K. https://livewarwickac.sharepoint.com/:v:/s/Flex-Covid19/EXfYLrNIaJNHpR-VJjjfbysB8yqrHSRHoTfSTeEjox4_XQ?e=84I094 (2020).
6. Roemer, R. A., Roemer, N. S. & Wallis, A. K. https://livewarwickac.sharepoint.com/:v:/s/Flex-Covid19/EUETBeGAatZOp4SGFew7P6ABPY1LSPsXNTm_ck2JjiGnaw?e=m4SRzh (2020).
7. Roemer, R. A., Roemer, N. S. & Wallis, A. K. https://livewarwickac.sharepoint.com/:v:/s/Flex-Covid19/EShhXMwZf8NMl2FdmQBARFsBxV9DbFR7K2bJpEMfys6Njg?e=sX5sbG (2020).
8. PanDDA analysis group. PanDDA analysis group deposition (2020).
9. Jin, Z. *et al.* Structure of M pro from COVID-19 virus and discovery of its inhibitors. *Nature* DOI: 10.1038/s41586-020-2223-y (2020).
10. Zhu, Y. & Sun, F. RCSB PDB - 6LVN: Structure of the 2019-nCoV HR2 Domain.
11. Xia, S. *et al.* Inhibition of SARS-CoV-2 (previously 2019-nCoV) infection by a highly potent pan-coronavirus fusion inhibitor targeting its spike protein that harbors a high capacity to mediate membrane fusion. *Cell Res.* **30**, 343–355, DOI: 10.1038/s41422-020-0305-x (2020).
12. Wang, Q. Q. *et al.* Structural and Functional Basis of SARS-CoV-2 Entry by Using Human ACE2. *Cell* **181**, 894–904, DOI: 10.1016/j.cell.2020.03.045 (2020).
13. Zhang, L. *et al.* Crystal structure of SARS-CoV-2 main protease provides a basis for design of improved a-ketoamide inhibitors. Tech. Rep. (2020). DOI: 10.1126/science.abb3405.
14. Lan, J. *et al.* Structure of the SARS-CoV-2 spike receptor-binding domain bound to the ACE2 receptor. *Nature* **581**, 215–220, DOI: 10.1038/s41586-020-2180-5 (2020).
15. Yan, R. *et al.* Structural basis for the recognition of SARS-CoV-2 by full-length human ACE2. *Science* **367**, 1444–1448, DOI: 10.1126/science.abb2762 (2020).
16. Su, H., Zhao, W., Li, M., Xie, H. & Xu, Y. RCSB PDB - 6M2N: SARS-CoV-2 3CL protease (3CL pro) in complex with a novel inhibitor.
17. Su, H., Zhao, W., Li, M., Xie, H. & Xu, Y. RCSB PDB - 6M2Q: SARS-CoV-2 3CL protease (3CL pro) apo structure (space group C21).
18. Chen, Y. *et al.* Biochemical and Structural Insights into the Mechanisms of SARS Coronavirus RNA Ribose 2'-O-Methylation by nsp16/nsp10 Protein Complex. *PLoS Pathog.* **7**, e1002294, DOI: 10.1371/journal.ppat.1002294 (2011).
19. Gao, Y. *et al.* Structure of the RNA-dependent RNA polymerase from COVID-19 virus. *Science* **368**, 779–782, DOI: 10.1126/science.abb7498 (2020).
20. Wrapp, D. *et al.* Cryo-EM structure of the 2019-nCoV spike in the prefusion conformation. *Science* **367**, 1260–1263, DOI: 10.1126/science.abb2507 (2020).
21. Shang, J. *et al.* Structural basis of receptor recognition by SARS-CoV-2. *Nature* **581**, 221–224, DOI: 10.1038/s41586-020-2179-y (2020).
22. Kim, Y. *et al.* Crystal structure of Nsp15 endoribonuclease <scp>NendoU</scp> from <scp>SARS-CoV</scp> -2. *Protein Sci.* pro.3873, DOI: 10.1002/pro.3873 (2020).
23. CSGID. CSGID analysis group (2020).
24. Walls, A. C. *et al.* Structure, Function, and Antigenicity of the SARS-CoV-2 Spike Glycoprotein. *Cell* **181**, 281–292, DOI: 10.1016/j.cell.2020.02.058 (2020).

**25.** Yuan, M. *et al.* A highly conserved cryptic epitope in the receptor binding domains of SARS-CoV-2 and SARS-CoV, DOI: 10.1126/science.abb7269 (2020).

**26.** Littler, D., Gully, B., Riboldi-Tunnicliffe, A. & Rossjohn, J. RCSB PDB - 6W9Q: Peptide-bound SARS-CoV-2 Nsp9 RNA-replicase.

**27.** Owen, C. *et al.* RCSB PDB - 6Y84: SARS-CoV-2 main protease with unliganded active site (2019-nCoV, coronavirus disease 2019, COVID-19).

**28.** Veverka, V. & Boura, E. RCSB PDB - 6YI3: The N-terminal RNA-binding domain of the SARS-CoV-2 nucleocapsid phosphoprotein.

**29.** Huo, J. *et al.* RCSB PDB - 6YLA: Crystal structure of the SARS-CoV-2 receptor binding domain in complex with CR3022 Fab.

**30.** Vuksanovic, N. & Silvaggi, N. RCSB PDB - 6WEY: High-resolution structure of the SARS-CoV-2 NSP3 Macro X domain.

**31.** Gupta, Y., Viswanathan, T., Arya, S. & Qi, S. RCSB PDB - 6WKS: Structure of SARS-CoV-2 nsp16/nsp10 ternary complex.

**32.** Kneller, D., Kovalevsky, A. & Coates, L. RCSB PDB - 6WQF: Structural Plasticity of the SARS-CoV-2 3CL Mpro Active Site Cavity Revealed by Room Temperature X-ray Crystallography.

**33.** Konkolova, E., Klima, M. & Boura, E. RCSB PDB - 6YHU: Crystal structure of the nsp7-nsp8 complex of SARS-CoV-2.

**34.** Huo, J. *et al.* RCSB PDB - 6YM0: Crystal structure of the SARS-CoV-2 receptor binding domain in complex with CR3022 Fab (crystal form 1).

**35.** Guenther, S. *et al.* RCSB PDB - 6YNQ: Structure of SARS-CoV-2 Main Protease bound to 2-Methyl-1-tetralone.

**36.** Huo, J. *et al.* RCSB PDB - 6YOR: Structure of the SARS-CoV-2 spike S1 protein in complex with CR3022 Fab.

**37.** Guenther, S. *et al.* RCSB PDB - 6YT8: Structure of SARS-CoV-2 Main Protease bound to pyrithione zinc.

**38.** Shin, D. & Dikic, I. RCSB PDB - 6YVA: PLpro-C111S with mISG15.

**39.** Ni, X. *et al.* Structural Genomics Consortium.

**40.** Hillen, H. *et al.* RCSB PDB - 6YYT: Structure of replicating SARS-CoV-2 polymerase.

**41.** Krafcikova, P., Silhan, J., Nencka, R. & Boura, E. RCSB PDB - 6YZ1: The crystal structure of SARS-CoV-2 nsp10-nsp16 methyltransferase complex with Sinefungin.

**42.** Fu, L. RCSB PDB - 7BRO: Crystal structure of the 2019-nCoV main protease.

**43.** Yin, W. *et al.* RCSB PDB - 7BV1: Cryo-EM structure of the apo nsp12-nsp7-nsp8 complex.

**44.** Wu, Y., Qi, J. & Gao, F. RCSB PDB - 7BZ5: Structure of COVID-19 virus spike receptor-binding domain complexed with a neutralizing antibody.

**45.** Pinto, D. *et al.* Seattle Structural Genomics Center for Infectious Disease.

**46.** Khan, M., Arutyunova, E., Young, H. & Lemieux, M. RCSB PDB - 6WTJ: Feline coronavirus drug inhibits the main protease of SARS-CoV-2 and blocks virus replication.

**47.** Sacco, M., Ma, C., Chen, Y. & Wang, J. RCSB PDB - 6WTT: Crystals Structure of the SARS-CoV-2 (COVID-19) main protease with inhibitor GC-376.

**48.** Lv, Z. & Olsen, S. RCSB PDB - 6WUU: Crystal structure of the SARS CoV-2 Papain-like protease in complex with peptide inhibitor VIR250.

**49.** Littler, D., Gully, B., Riboldi-Tunnicliffe, A. & Rossjohn, J. RCSB PDB - 6WXD: SARS-CoV-2 Nsp9 RNA-replicase.

**50.** Ye, Q. & Corbett, K. RCSB PDB - 6WZO: Structure of SARS-CoV-2 Nucleocapsid dimerization domain, P1 form.

**51.** Henderson, R. & Acharya, P. RCSB PDB - 6X29: SARS-CoV-2 rS2d Down State Spike Protein Trimer.

**52.** Zinzula, L., Basquin, J., Nagy, I. & Bracher, A. RCSB PDB - 6YUN: 1.45 Angstrom Resolution Crystal Structure of C-terminal Dimerization Domain of Nucleocapsid Phosphoprotein from SARS-CoV-2.

**53.** Peng, Q., Peng, R. & Shi, Y. RCSB PDB - 7BW4: Structure of the RNA-dependent RNA polymerase from SARS-CoV-2.

**54.** Shi, R., Qi, J., Wang, Q., Gao, F. & Yan, J. RCSB PDB - 7C01: Molecular basis for a potent human neutralizing antibody targeting SARS-CoV-2 RBD.

**55.** Zhou, R., Zeng, R. & Lei, J. RCSB PDB - 7C22: Crystal structure of the C-terminal domain of SARS-CoV-2 nucleocapsid protein.

**56.** Lin, S. *et al.* RCSB PDB - 7C2I: Crystal structure of nsp16-nsp10 heterodimer from SARS-CoV-2 in complex with SAM (with additional SAM during crystallization).
